# Genome-wide analysis of the role of the antibiotic biosynthesis regulator AbsA2 in *Streptomyces coelicolor* A3(2)

**DOI:** 10.1101/361360

**Authors:** Richard A. Lewis, Abdul Wahab, Giselda Bucca, Emma E. Laing, Carla Möller-Levet, Andrze Kierzek, Colin P. Smith

**Author notes:** Corresponding authors (RAL); (CPS).

## Abstract

The AbsA1-AbsA2 two component signalling system of *Streptomyces coelicolor* has long been known to exert a powerful negative influence on the production of the antibiotics actinorhodin, undecylprodiginine and the Calcium-Dependent Antibiotic (CDA). Here we report the analysis of a *ΔabsA2* deletion strain, which exhibits the classic precocious antibiotic hyper-production phenotype, and its complementation by an N-terminal triple-FLAG-tagged version of AbsA2. The complemented and non-complemented *ΔabsA2* mutant strains were used in large-scale microarray-based time-course experiments to investigate the effect of deleting *absA2* on gene expression and to identify the *in vivo* AbsA2 DNA-binding target sites using ChIP-on chip. We show that in addition to binding to the promoter regions of *redZ* and *actII*-*orfIV* AbsA2 binds to several previously unidentified sites within the *cda* biosynthetic gene cluster within and/or upstream of *SCO3215* - *SCO3216*, *SCO3217*, *SCO3229* - *SCO3230*, and *SCO3226*, and we relate the pattern of AbsA2 binding to the results of the transcriptomic study and antibiotic phenotypic assays. Interestingly, dual ‘biphasic’ ChIP peaks were observed with AbsA2 binding across the regulatory genes *actII*-*orfIV* and *redZ* and the *absA2* gene itself, while more conventional single promoter-proximal peaks were seen at the CDA biosynthetic genes suggesting a different mechanism of regulation of the former loci. Taken together the results shed light on the complex mechanism of regulation of antibiotic biosynthesis in *Streptomyces coelicolor* and the important role of AbsA2 in controlling the expression of three antibiotic biosynthetic gene clusters.

## Introduction

The bacteria of the genus *Streptomyces* are notable for their ability to undergo morphological differentiation and for their ability to synthesize a wide variety of secondary metabolites. The model streptomycete, *Streptomyces coelicolor* A3(2), produces several antibiotics, including the non-ribosomally synthesised lipopeptide Calcium Dependent Antibiotic, CDA [1], the blue pigmented polyketide, actinorhodin (ACT) [2] and the red pigmented undecylprodiginines (RED) [3]. The genes responsible for the biosynthesis of each of the antibiotics are located in distinct clusters [2-4] each of which comprise a SARP (streptomycete antibiotic regulatory protein) gene encoding a pathway specific activator protein *i*.*e*. *cdaR* [5], *redZ* [6], *redD*, [7] and *actII*-*orfIV*, [8-9]. Induction of antibiotic biosynthesis is closely linked to morphological differentiation and the transcription of genes involved in both processes are controlled by complex regulatory networks in response to environmental and physiological stimuli [10]. The *absA* locus was originally identified by isolating mutants unable to synthesize antibiotics but unaffected in their morphological development [11]. Sequencing of this locus identified two genes, *absA1* and *absA2* which comprise a two-component system, with AbsA1 possessing similarity to other streptomycete antibiotic biosynthetic cluster associated histidine-kinase sensor-transmitter proteins and AbsA2 displaying similarity to DNA-binding response regulator proteins [12]. The location of *absA1/2* within the CDA biosynthetic gene cluster was only demonstrated after the determination of the *S*. *coelicolor* genome sequence [5, 13-14].

Sequencing of the original *absA* mutants by the Champness laboratory, together with suppressor mutants [15], and introduction of a series of deletion and point mutations [16], strongly indicated that the active, autophosphorylated form of AbsA1 was responsible for generating the repressive, phosphorylated form of AbsA2. Biochemical experiments investigating the activity of AbsA1 have confirmed the results of the genetic studies and showed that AbsA1 is able to phosphorylate and dephosphorylate AbsA2 [17]. The effect of gene dosage on the activity of AbsA1 & 2 was also investigated and the fact that the *absA1/2* transcript was more abundant in the original *absA1* mutant C542 [11] which possesses two amino acid changes in a region thought to be involved in aspartylphosphatase activity [15] and less abundant in the *absA2* mutant C570, in which the phosphorylated aspartate residue has been changed to a glutamate residue [16], relative to the parent strain J1501, suggested that *absA2* is positively autoregulated and that the autoregulatory form is phosphor-AbsA2 [16].

S1 nuclease transcript mapping indicated that *absA1* and *absA2* were expressed as a single bi-cistronic leaderless transcript from the promoter upstream of *absA1* (P1) [16]. However, *absA2* is also expressed from a second promoter (P2) located inside *absA1* [18], which is is 4-7 times stronger than that of P1, based on transcript abundance and reporter assays and the expression of *absA2* is consequently higher than that of *absA1*. Significantly, the relative abundances of the two transcripts appear not to vary throughout growth or with growth media type [18]

Ryding *et al*., [19] investigated the effect on the expression of the genes of the *cda* cluster in J1501 and its derivative mutant *absA* C542 and detected expression from the promoters upstream of *SCO3230* (*cdaPSI*), *SCO3249* (*acp*), *SCO3229* (*hmaS*) and *SCO3215* (*glmT*) in J1501 but not in the C542 mutant. Expression from the promoters upstream of *SCO3225* (*absA1*) and *SCO3224* (a putative CDA resistance gene), was detected in C542 but not in J1501. Significantly, this study revealed little evidence for the regulation of *cdaR* by AbsA1/2 although more conclusive evidence was found for the AbsA1/2 mediated regulation of *redZ* [19], and the involvement of the *absA* locus in regulating *actII*-*orfIV* and *redD* has also been noted [20]. Gel retardation assays, using phosphorylated and non-phosphorylated tagged versions of AbsA2, to investigate its binding to the promoters of *SCO3217* (*cdaR*), *SCO3226*, (*absA1/2*) and *SCO3230* (*cdaPSI*) were unable to show that either form of AbsA2 bound the respective promoter regions, which suggested that *in vivo* AbsA2 may bind cooperatively with another protein *e*.*g*. CdaR [17] or require a particular DNA topology. More recently ChIP studies using anti-AbsA2 antibodies to pull-down AbsA2 complexes were conducted, however, the only genes whose promoters were found to be bound by AbsA2 were those of *cdaR*, *redZ* and *actII-orfIV* [21].

The results of McKenzie & Nodwell, (2007) suggest AbsA2 plays a role as a “master” regulator of antibiotic biosynthesis and regulates the SARPs for the *cda*, *act* and *red* gene clusters [21]. The present large-scale, time-course transcriptomic experiment, conducted in parallel with an *in vivo* AbsA2 ChIP-on-chip study, was conceived to more comprehensively examine the role of AbsA2, particularly with respect to the CDA biosynthetic gene cluster where several promoters have been previously shown to be AbsA2 dependent [19].

## Materials & Methods

### Strains

The *Streptomyces coelicolor* strain used in this study was MT1110 [22] which is a prototrophic SCP1-, SCP2-derivative of the wild-type strain 1147 [23]. The construction of the MT1110 *ΔcdaR* mutant is provided in [24] where the deletion of the *EcoRI* fragment (+442 to +1174 bp) is described. Plasmid maintenance was done in *E*. *coli* JM109 (*endA1*, *rec*A1, *gyr*A96, *thi*-1, *hsd*R17, (r_k_^-^ m_k_^+^), *mcr*A, *rel*A1, *sup*E44λ^-^, Δ(*lac pro*B), [F’ *tra*D36, *pro*AB+, *lac*I^q^,Δ(*lac*Z)M15). The *E*. *coli* strain ET12567 (F^-^ *dam*-*13*::Tn*9 dcm*-*6 hsdM hsdR recF143 zjj*-*202*::Tn*10 rspL136*) [25] was used for propagation of plasmid DNA free from *dam*, *dcm* and *hsd* methylation and when containing the conjugative vector pUB307, was used in *E*. *coli/S*. *coelicolor* conjugations [26].

### Construction of a *ΔabsA2* strain and *absA2*-complementing plasmid

An *absA2* deletion mutant was constructed using homologous recombination to replace the wild-type gene with a truncated version containing an internal deletion of 618 bp out of a total of 669 bp. An *Xba*I site was substituted for the wild-type nucleotides between positions 3538686 and 3539309 [14], so introducing a termination codon into the truncated version of *absA2*, which terminates translation after incorporation of three amino acids (MIL). A 1,757 bp fragment containing part of the *absA2* N-terminal coding sequence and upstream region was amplified by PCR using primers “absA2up forward” and “absA2up reverse” (Table 1) and cloned into pKC1132 as an *Eco*RI-*Xba*I fragment to generate pAW3. A 1,503 bp fragment containing the C-terminal coding sequence of *absA2* and part of the downstream region was amplified by PCR using primers “absA2down forward” and “absA2down reverse” (Table 1) and cloned as an *Xba*I-*Hind*III fragment into pAW3 to generate pAW33. Plasmid DNA of pAW33, obtained from ET12567, was used to transform MT1110 protoplasts [23] and primary apramycin resistant transformants were selected. Following three rounds of non-selective growth Apra^S^ colonies were identified by replica plating and the presence of the *ΔabsA2* truncation assayed for by PCR, followed by confirmation by Southern blotting. The mutant strain was designated MT1110 *ΔabsA2*.

**Table 1.**
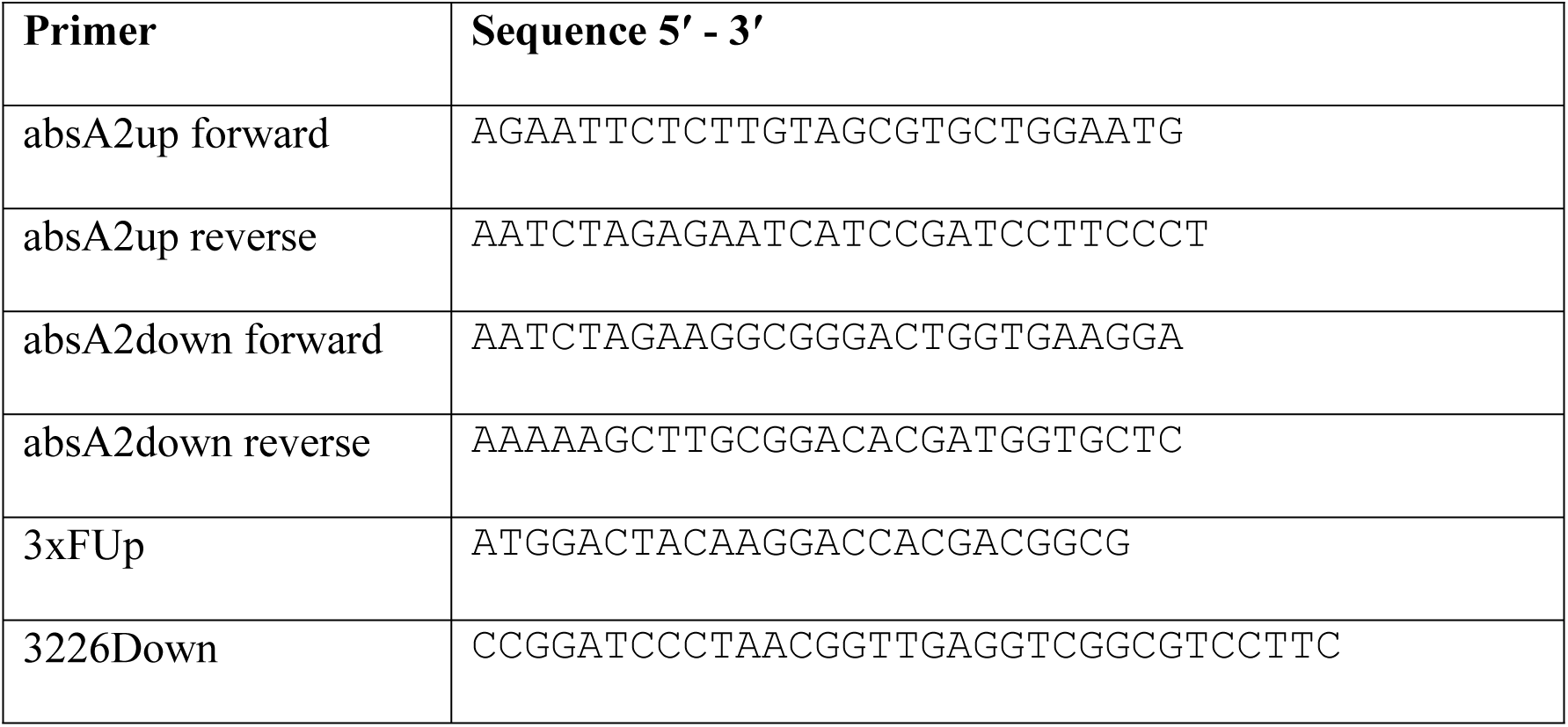
List of DNA sequences of named oligonucleotides used in the present study.

For the construction of a *ΔabsA2* complementing plasmid an N-terminally triple-FLAG-tagged version of *absA2* was synthesized *de novo* by GenScript^™^ and supplied cloned into the *Eco*RV site of pUC57. The 773 bp insert (Fig. S1) comprised the *absA2* coding sequence, fused at the N-terminus to a sequence encoding a triple-FLAG tag (3xF), together with the *absA1*-*absA2* intergenic region and the *absA2* ribosome binding site. This insert does not comprise the P2 promoter [18]. The insert was sub-cloned as a *Bam*HI - *Xba*I fragment into similarly cut pMT3226 [23], so that *absA2* was under the control of the tandem glycerol-inducible (*gyl*) promoters. The resultant plasmid pMT3226::*3xF*-*absA2* was used to transform ET12567 (pUB307). This construct, and the parent vector pMT3226 as a negative control, were introduced into MT1110 *ΔabsA2* by conjugation [26]. The presence of the 3xF tagged version of *absA2* was confirmed in Apra^R^ ex-conjugants by PCR using primers “3xFUp” and “3226Down”. The pigmented antibiotic phenotypes of the complemented mutant (MT1110 *ΔabsA2* (pMT3226::*3xF*-*absA2*)) and non-complemented mutant (MT1110 *ΔabsA2* (pMT3226)) strains were determined visually following growth on solid media plates *i*.*e*. Mannitol-Soya agar [23] and Oxoid^™^ Nutrient Agar (ONA) (Oxoid^™^), supplemented with 0.5% glycerol, to induce expression from the *gyl* promoters.

### Bacterial culture

Spores were produced from confluent lawns of *Streptomyces coelicolor* cultures grown on Mannitol-Soya agar [23] and were pre-germinated by incubating at 30°C, with shaking (200 r.p.m) in 2YT (with the addition of 30% sucrose) for eight hours. Duplicate cultures for both the complemented (MT1110 *ΔabsA2* (pMT3226::*3xF*-*absA2*)) and *ΔabsA2* (MT1110 *ΔabsA2* (pMT3226)) strains were set up in parallel. Pre-germinated spores were used to inoculate two-litre spring flasks containing 500 ml of modified YEME [23] to give an O.D._450_ of 0.06. The modified YEME comprised 10% sucrose instead of 34% and glucose was replaced by glycerol. Apramycin (50 μg/ml) and 0.01% Antifoam 204 (Sigma^™^) were also added. The cultures were incubated at 30°C, with shaking (200 r.p.m) and growth was monitored by following the O.D._450_. Samples were taken for transcriptomic, ChIP-on-chip and pigmented antibiotic assays at 14, 18 and 35 h after inoculation.

### Culture sampling and antibiotic assays

For transcriptomic studies 10 ml samples of culture were harvested as described in [27]. For ChIP-on-chip studies 108 ml of each culture was formaldehyde cross-linked and quenched as described by Allenby *et al*. [28] and divided into three 42 ml portions before being pelleted (4,000 r.p.m, 4°C, 10 min), and the supernatants discarded. The pellets were each washed twice in 20 ml of PBS and stored at −20°C. For pigmented antibiotic assays 10 ml of whole culture was frozen at −20°C. CDA bioassays were conducted according to the method of Chong *et al*., [13]. Actinorhodin assays were conducted according to the method of Bystrykh *et al*., [29] modified as described by Lewis *et al*., [27], whilst undecylprodiginine assays were conducted according to the method of Tsao *et al*., [30], modified as described by Lewis *et al*., [27].

### Nucleic acid isolation and ChIP-on-chip techniques

For this study a protocol, based on the standard protocol “Total RNA extraction by Tissue Lyser”, described previously [31],was developed which allows the isolation of all RNA species, including small RNAs. Following the passaging of the sample through the Qiagen^™^ RNeasy^™^ gDNA eliminator column, the flow-though was mixed with an appropriate volume of ethanol and used in the protocol “F.1. Total RNA Isolation Procedure” according to the Ambion^™^ “*mir*Vana^™^ miRNA Isolation Kit” instructions. The quality of the RNA obtained was assessed by running a Bioanalyzer Prokaryote Total RNA Nano chip (Agilent^™^) and only RNA having a RNA Integrity Number of 6.5 or above was used in subsequent work. Protocols for *S*. *coelicolor* MT1110 genomic DNA isolation and labelling (Cy5), together with the labelling of cDNA (Cy3) were as previously described [28] and hybridizations were set up as before [32] using 2 x 105 K high-density IJISS *Streptomyces* whole genome microarrays [33]. The transcriptomic microarray data are deposited with ArrayExpress (Accession Number:-E-MTAB-3528).

Protocols for the chromatin extraction and immunoprecipitation techniques for the ChIP-on-chip experiment were as previously described [32] with the modifications that the antibody used was the M2 mouse monoclonal anti-FLAG antibody from Sigma^™^ and the pull-down was effected using Protein G magnetic beads (NEB^™^). The DNA obtained was labelled and hybridised to 2 x 105 K *Streptomyces* microarrays, as previously described by Lewis *et al*., [33]. The antibody immunoprecipitated chromatin was co-hybridized directly with mock “no antibody” immunoprecipitated chromatin *i*.*e*. the sample processed in the same way as the antibody immunoprecipitated chromatin, but without use of the specific antibody. To compensate for any dye bias in the experiment replicate hybridisations were conducted on different arrays with chromatin samples labelled in opposite Cy-dye orientations. The ChIP-on-chip microarray data are deposited with ArrayExpress (Accession Number:-E-MTAB-3527).

### Microarray chIP data processing

#### Pre-processing

Microarray image files were scanned and imported into the Agilent Feature Extraction software (v9.1) for image analysis and resulting files were processed in R. Log_2_ values were scale-normalized using the “normalizeMedianAbsValues” function in R Bioconductor package Limma [34-35]. Probes were filtered out based on Agilent feature flags QC metrics [36]. In each time point, probes were excluded if they were flagged (in both channels) in one or more samples in both strains or in more than two samples.

### AbsA2 enrichment and regions of binding

Candidate probes for AbsA2 enrichment were defined as those for which their value in the complemented mutant (3xF) sample is larger than their value in the non-complemented (*ΔabsA2*) mutant sample in both replicates. Candidate probes with a fold-change (defined as the difference between the averaged 3xF values and averaged *ΔabsA2* values) larger than 1.25 standard deviations from the mean fold-change across all candidate probes were defined as AbsA2-enriched probes. These probes were then clustered based on their genomic position and 1 kb regions in which at least two enriched probes were identified were considered to be regions of binding, and thus of interest.

### Microarray expression data processing

As for the ChIP-on-chip data, the raw expression data Feature Extraction files were imported into R for processing with Limma. No spatial effects were found, therefore all arrays within the experiment were normalised by the ‘global median within array’, followed by the ‘scale across array’ normalisation methods. Poor quality probes, as determined by the Agilent software, were filtered from the data-set. The representative expression signal for a gene was calculated by averaging all good quality probes that target the annotated coding region for that gene. Thus, for each time-point in each condition two values, one for each biological replicate, were obtained for every gene in the *S*. *coelicolor* genome. The replicated gene data were analysed for significant differential expression between the complemented and non-complemented strains using Rank Products analysis *via* the web-tool RankProdIt [37-38].

## Results & discussion

### Antibiotic phenotypes of the complemented and non-complemented *ΔabsA2* strains

The deletion from *S*. *coelicolor* MT1110 of the majority of the *absA2* coding sequence coupled with the introduction of a termination codon was expected to confer the classic precocious hyperproduction of antibiotic (*pha*) antibiotic phenotype on MT1110 with regard to CDA, ACT and RED biosynthesis, [16]. The non-complemented mutant strain (MT1110 *ΔabsA2* (pMT3226) over-expressed ACT and RED relative to the wild-type strain on a number of different media, whilst the complemented mutant (MT1110 *ΔabsA2* (pMT3226::*3xF*-*absA2*)) exhibited a hyper-repressed pigmented antibiotic phenotype (Fig. 1(A) & Fig. 1(B)). The repression of ACT and RED production by introduction of the heterologously expressed version of *absA2* demonstrates that the N-terminally triple-flag tagged version of AbsA2 is capable of complementing the *ΔabsA2* and is likely to be a substrate for AbsA1 kinase activity, given that phosphorylated AbsA2 is known to be the functional form of the protein, and it is presumably capable of binding to the *actII*-*orfIV* and *redZ* promoters.

**Fig. 1.**
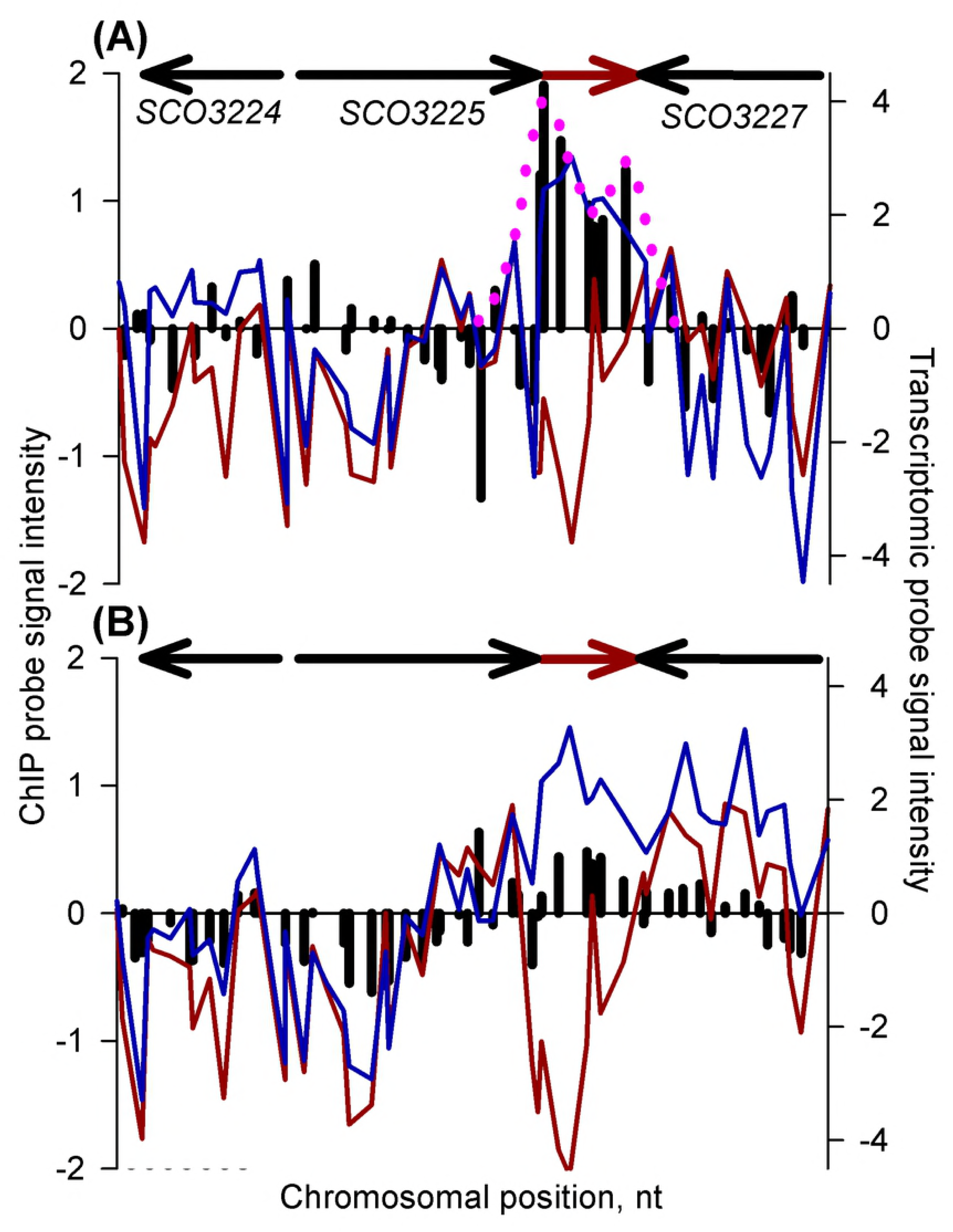
Pigmented antibiotic phenotypes of strains used in the study.

MS agar (A) and ONA (B) plates illustrating the ACT and RED phenotypes of: (a) MT1110; (b) MT1110 *ΔabsA2* (pMT3226); (c) MT1110 *ΔabsA2* (pMT3226::*3xFabsA2*)

Surprisingly the CDA over-production phenotype, (Fig. 2(A) & Fig. 2(B)) seen in MT1110 *ΔabsA2* (pMT3226) was not reversed by introduction of the N-terminally triple-FLAG tagged version of *absA2*. We will return to discuss this intriguing phenomenon below.

**Fig. 2.**
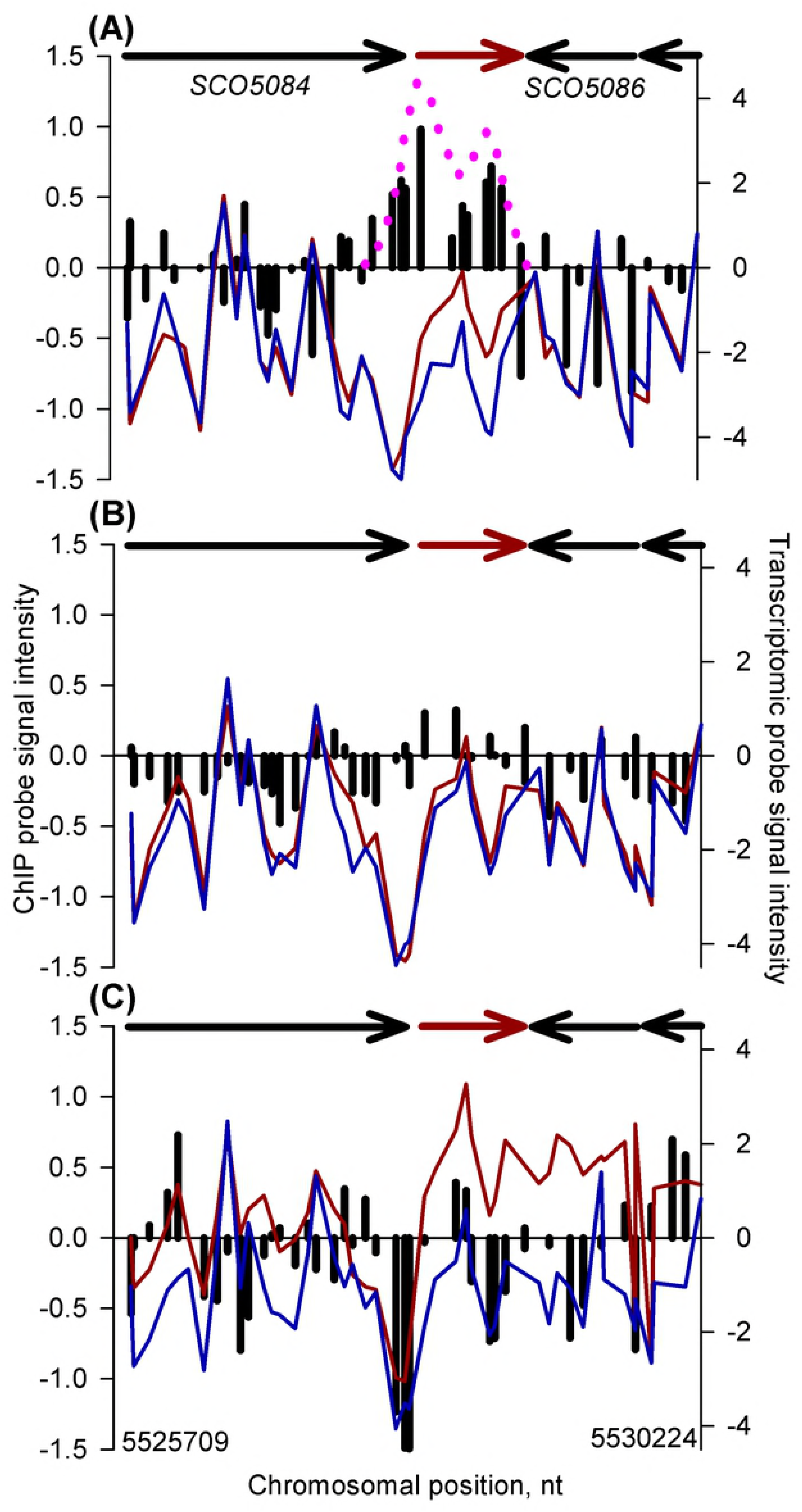
Calcium dependent antibiotic phenotypes of strains used in the study.

CDA bioassay plates, plus Ca^2+^ (A) and minus Ca^2+^ (B) illustrating the CDA production phenotypes of: (a) MT1110; (b) MT1110 *ΔabsA2* (pMT3226); (c) MT1110 *ΔabsA2* (pMT3226::*3xFabsA2*); (d) MT1110 *Δcda*.

The time points at which the liquid cultures were sampled were chosen to represent mid-exponential phase (14 h) *i*.*e*. before the onset of pigmented antibiotic production, late exponential phase (18 h) which correlated with the onset of RED biosynthesis (as determined by visual inspection) and stationary phase (35 h) when ACT biosynthesis was well advanced. The results of the ACT assays on the liquid cultures, (Fig. S2) are consistent with the antibiotic phenotypes observed on solid media with the *ΔabsA* strain producing actinorhodin whilst the *3xFabsA* strain produced no detectable ACT.

### Transcriptomic analysis of the complemented and non-complemented *AabsA2* strains

The results (Fig. S3) of the *absA2* transcriptomic study indicated that, as expected from the results of the antibiotic phenotypic assays, a large number of antibiotic biosynthetic genes were significantly differentially expressed in the *3xFabsA2* strain relative to the *ΔabsA2* strain.

At stationary phase (35 h) sixteen genes of the actinorhodin biosynthetic gene cluster are up-regulated in the *ΔabsA2* strain relative to the *3xFabsA* strain which correlates well with the results of the ACT assays in liquid culture, (Fig. S2) and the phenotypic assays on solid media, (Fig. 1(A) & Fig. 1(B)).

The fact that no genes of the *red* biosynthetic gene cluster are represented amongst the significantly differentially expressed genes was expected given the similarity of the RED phenotypes of both the *3xFabsA2* and *ΔabsA2* strains in liquid culture, (Fig. S2).

The results of the CDA bioassay are consistent with the results of the transcriptomics experiment which demonstrate that many genes of the *cda* cluster are well represented in the lists of significantly differentially expressed genes, being up-regulated in the complemented mutant relative to the non-complemented mutant. Three genes of the *cda* cluster are up-regulated in the *3xFabsA2* strain relative to the *ΔabsA2* strain at the mid exponential time-point (14 h), whilst nineteen *cda* genes are up-regulated in the *3xFabsA2* strain relative to the *ΔabsA2* strain at the late-exponential phase time-point (18 h) (Fig. S3). Consideration of these results must of course exclude *SCO3226* (*absA2*) whose differential expression was expected due to its deletion from the *ΔabsA2* strain, rather than due to *bona fide* changes in its expression.

When the results of the transcriptomic study are compared with the AbsA2 “network module” of Castro-Melchor *et al*., [39] with the exceptions of *SCO3220*-*3224* (*cda* genes) none of these genes appear in the list of significantly differentially expressed genes. If indeed the expression of these “network module” genes is truly linked to AbsA2 then they are not directly, nor indirectly, regulated by AbsA2 under the conditions of the present study, nor are they significantly differentially expressed concomitantly.

The same study [39] also described a CdaR “network module”. However, these genes do not correlate with the results presented herein, excepting *SCO7717* which is up-regulated in the complemented *ΔabsA2* strain relative to the non-complemented *ΔabsA2* strain at the 18 h time-point. As *cdaR* is similarly up-regulated at this time-point this is consistent with CdaR being a positive regulator of *SCO7717*.

Although the function of *SCO7717* is not known a similarity search indicates that the gene most similar to it in *S*. *coelicolor* is the *cda* cluster located gene, *SCO3220* whose function is also unclear. It is well known that there is cross-talk/redundancy between gene products encoded by secondary metabolite clusters in that proteins encoded by one cluster are capable of substituting for similar proteins in other cluster. For example, the NRPS adenylation domain activation protein, CdaX, (SCO3218) of the *cda* cluster is able to substitute for its counterpart, CchK, in the coelichelin biosynthetic cluster [40]. It is therefore possible that SCO3220 and SCO7717 are functionally redundant and that SCO7717 has usurped the role of SCO3220 in CDA biosynthesis and its gene is thus co-regulated with the *cda* cluster genes by CdaR.

### ChIP-on-chip analysis of AbsA2 in the FLAG-tagged *absA2* and *ΔabsA2* strains

To investigate whether the significantly differentially expressed genes (Fig. S3) were directly regulated by AbsA2 we conducted ChIP-on-chip studies on samples taken at the same time-points as those for the transcriptomic studies. The genes with background shading in Fig. S3 are those which also feature in the ChIP-on-chip enriched probe cluster list. It is difficult to interpret the majority of these in terms of the known function of AbsA2 *i*.*e*. regulation of antibiotic biosynthesis. Whilst we cannot exclude the possibility that AbsA2 does regulate these genes, an in depth visual analysis of the chIP binding data, in combination with the expression patterns, and knowledge of gene function, has been unable to establish meaningful correlations between AbsA2 binding and gene expression/function for the majority of these genes. We therefore confine further discussion/analysis of the results to known, validated AbsA2 targets (*i*.*e*. *cdaR*, *redZ* and *actII*-*orfIV*) and to genes of the *cda* cluster, based on the results of a previous *absA2* deletion study [19].

Although the transcriptomic results indicate AbsA2 exerts a strong effect on expression of the *cda* genes it was unclear whether these effects were directly, or indirectly, mediated by AbsA2, therefore, the results of the ChIP study relating to the *cda* cluster were of particular interest. Indeed, the ChIP data indicated that AbsA2 does bind to several sites within the *cda* cluster, binding to a region upstream of *SCO3217* (*cdaR*), to the intergenic region of *SCO3215* (*glmT*) and *SCO3216* (*ATPase*), to the intergenic region between *SCO3229* (*hmaS*) and *SCO3230* (*cdaPSI*), and lastly to another region within *SCO3226* (*absA2*).

The AbsA2 ChIP peak which can most obviously be linked to a change in expression of a particular gene at a certain time-point is the stationary phase peak seen in the *SCO3215*-*SCO3216* intergenic region which comprises two divergent promoters (Fig. 3). This region does not contain ChIP peaks at the mid and late exponential phase time-points and the appearance of the stationary phase peak correlates with a decrease in expression of *SCO3215* in the *3xFabsA2* strain, relative to its level at the late exponential phase time-point. The expression levels of the flanking genes remain similar in both strains at all time-points. The biological significance of the stationary phase repression of *SCO3215* (*glmT*), and presumably a consequential decrease in production of forms of CDA comprising methyl-glutamate, is unclear.

**Fig. 3.**
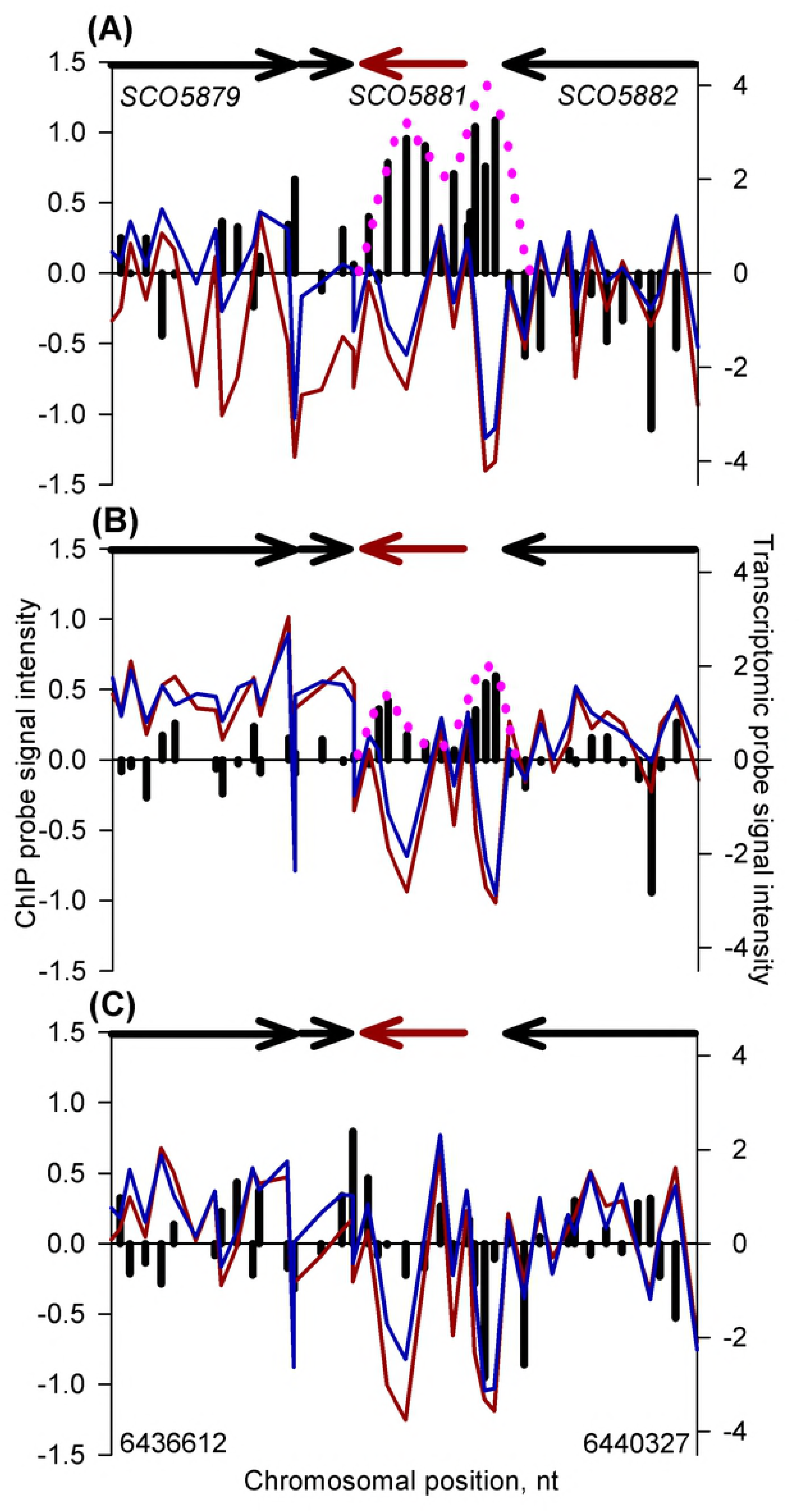
*In vivo* genomic distribution of 3xFAbsA2 in the region of *glmT* (*SCO3215)* and parallel measurement of adjacent gene expression.

The right hand *y* axis represents transcriptomic data (average log_2_ [cDNA/gDNA]) for MT1110 *ΔabsA2* (pMT3226) (red line) and MT1110 *ΔabsA2* (pMT3226::*3xFabsA2*) (blue line). The left hand *y* axis represents chIP data (average log_2_ [complemented mutant/non-complemented mutant enrichment signal]) [*i*.*e*. the Cy-dye balanced average MT1110 *ΔabsA2* (pMT3226::*3xFabsA*) enrichment ratio relative to the same probe signal from the same chromatin subjected to a mock no-Ab IP **divided** by the Cy-dye balanced average MT1110 *ΔabsA2* (pMT3226) enrichment ratio relative to the same probe signal from the same chromatin subjected to a mock no-Ab IP] (plotted as black columns). The *x* axis represents the genomic position of the microarray probes and the arrows above represent genes and direction of transcription, with the main gene of interest being highlighted in red. Panels (A), (B) & (C) refer to the 14 h (mid exponential phase), 18 h (late exponential phase) and 35 h (stationary phase) time-points respectively.

The relationship between AbsA2 binding and gene expression appears more complex at the *SCO3229* (*hmaS*) – *SCO3230* (*cdaPSI*) - intergenic region. Although there are AbsA2 ChIP peaks present at this locus at both the 14 and 18 h time-points in the *3xFabsA2* strain (Fig.4), there is little difference in expression of the two genes between the *3xFabsA2* and *ΔabsA2* strains at the 14 h time-point, although expression of both is markedly elevated in the *3xFabsA2* strain relative to the *ΔabsA2* strain at the 18 h time-point. At the 35 h time-point no ChIP peak is present and *hmaS* and *cdaPSI* are similarly expressed in both strains. This result is intriguing since, given the wealth of previous genetic studies relating to AbsA2, it was expected that an AbsA2 binding event would correlate with *repression* of transcription, as a previous study reported that the C542 *absA* mutant repressed expression from both the *SCO3230* and *SCO3229* promoters, relative to the parent strain, [19]. The lack of striking correlation between AbsA2 binding events and expression levels suggests that in addition to AbsA2 some other factor, or factors, is/are also involved in the regulation of *cdaPSI* and it is very likely that CdaR plays an activatory role here.

**Fig. 4.**
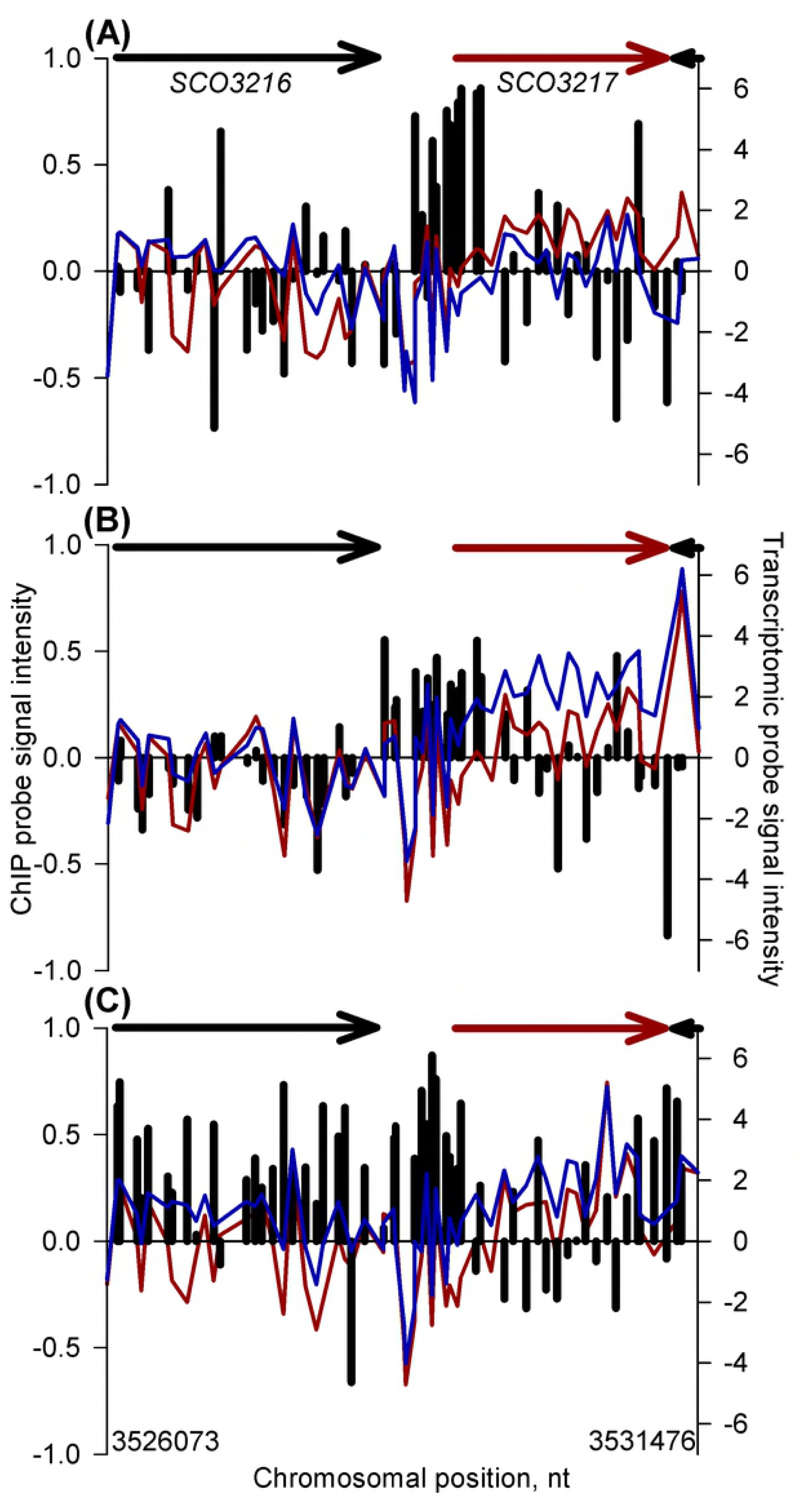
*In vivo* genomic distribution of 3xFAbsA2 in the region of *cdaPSI* (*SCO3230)* and parallel measurement of adjacent gene expression.

See legend to Fig. 3 for an explanation of the axes and colour coding. Panels (A), (B) & (C) refer to the 14 h, 18 h and 35 h time-points respectively.

The presence of a strong ChIP peak upstream of *cdaR* (SCO3217)(Fig. 5) was expected given the results of a previous ChIP study, [21]. AbsA2 peaks are observed at all three time-points, (specific binding is most clearly demonstrated at the 14 and 18 h time-points) although the effect of AbsA2 binding does not correlate with *cdaR* transcriptional activity, as expression of *cdaR* is slightly repressed in the *3xFabsA2* strain relative to the *ΔabsA2* strain at the 14 h time-point, but then is more highly expressed at the 18 h and 35 h time-points. AbsA2 binding *per se* may not be sufficient to regulate *cdaR* expression; for example, CdaR is known to be autoregulatory (A. E. Hayes, Z. Hojati and CPS, unpublished data).

**Fig. 5.**
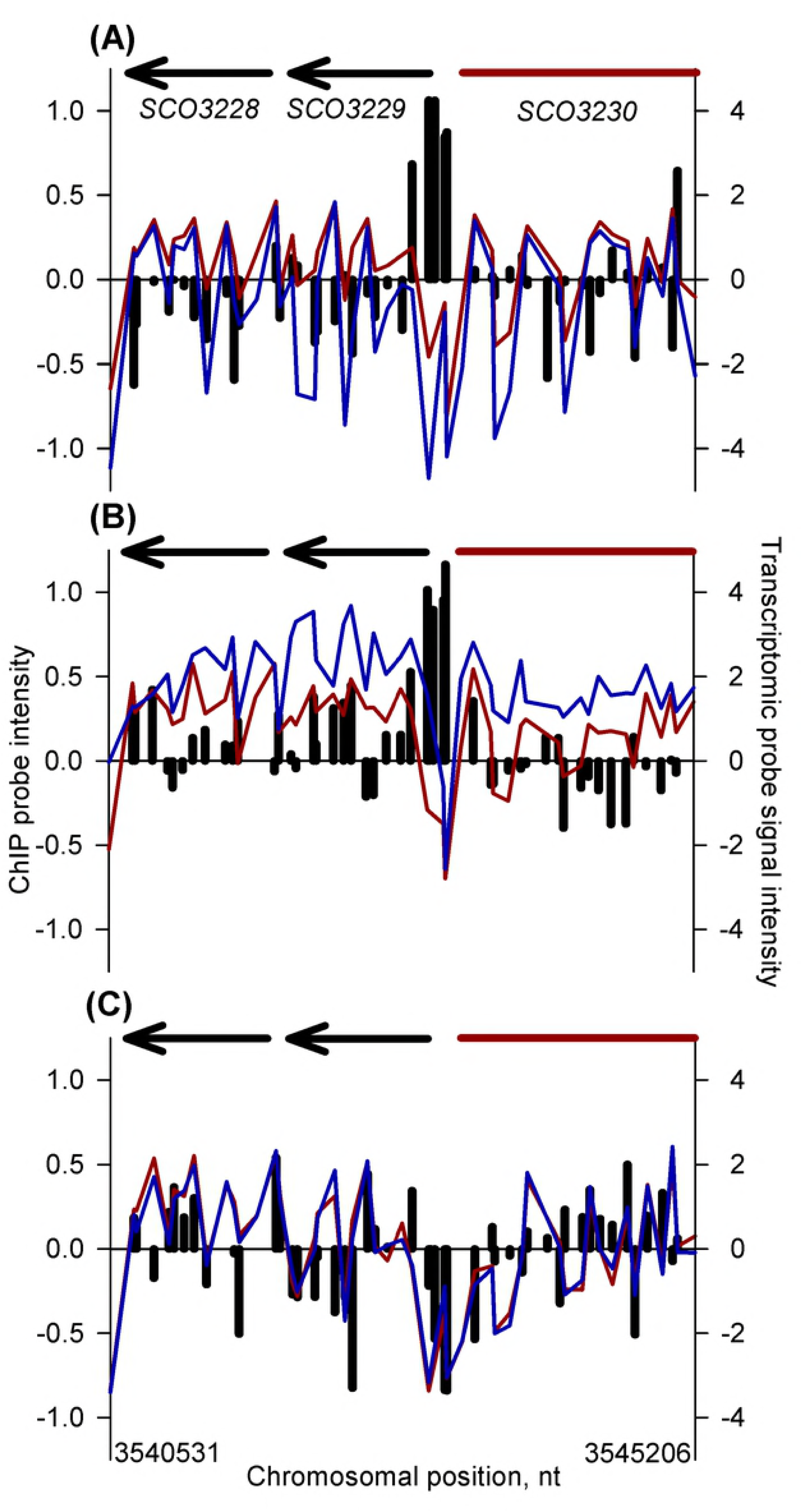
*In vivo* genomic distribution of 3xFAbsA2 in the region of *cdaR* (*SCO3217)* and parallel measurement of adjacent gene expression.

See legend to Fig. 3 for an explanation of the axes and colour coding. Panels (A), (B) & (C) refer to the 14 h, 18 h and 35 h time-points respectively.

The situation observed here with regard to *SCO3230* and *SCO3217* is consistent with the regulatory model proposed by Ryding *et al*., [19] who suggested “*that the AbsA2 protein directly regulates the cda biosynthetic promoters and in its negatively regulating form overrides any positive regulation that may be exerted by cdaR*” and they go on to suggest that *“the negative regulatory effect of AbsA2 on CDA synthesis may serve to modulate cda expression in the wild-type, perhaps in competition with cdaR putative activation*”.

We believe that at the *cdaR* and *cdaPSI* promoters the triple-flag tagged version of AbsA2 present in the complemented mutant is able to bind DNA in a normal wild-type fashion giving rise to the AbsA2 ChIP peaks observed. However, we suggest that unlike the wild-type AbsA2 the tagged version of the protein is unable to interact with, and repress, the transcription-activating effect of CdaR presumably due to the presence of the N-terminal triple-flag tag. Therefore, although we see clear evidence of AbsA2 binding at these promoters we do not see AbsA2 mediated transcriptional repression, as presumably CdaR is still active. This model satisfactorily explains the non-complementation of the *pha* phenotype with regard to CDA by the triple-flag tagged version of AbsA2 (Fig. 2(A) & Fig. 2(B)). It is likely that the hyper-production of CDA by the *3xFabsA2* strain, together with the enhanced expression of *cdaR* and *cdaPSI* in the complemented mutant strain at the mid-exponential time-point (Fig. 4 & Fig. 5) is due to the significant up-regulation of *cdaR* in the *3xFabsA2* strain at the mid-exponential phase time-point, combined with the inability of the 3xF tagged version of AbsA2 to repress the activating effect of CdaR. The apparent AbsA2-mediated repression seen at the *glmT* promoter (Fig. 3) may be a genuine instance of AbsA2 directly mediating repression, or may merely reflect a reduction in CdaR activity at this promoter during stationary phase.

The existence of a modified form of *absA2* which possesses differential activities with regard to CDA and ACT and RED biosynthesis is not unprecedented. When screening for mutations able to suppress *absA* mutations Anderson *et al*., [15], discovered a mutant, C577S20, which possessed a mutation encoding a V29A change in the N-terminus of AbsA2 and which produced ACT and RED but not CDA [15]. Although this situation is the opposite to that described herein it does suggest that AbsA2 operates through two different repressive mechanisms, one specific for CDA and the other for ACT and RED.

The final AbsA2 ChIP peak which targets the *cda* cluster is that which targets the internal region of the *absA2* gene (*SCO3226)* at the mid exponential phase time-point, (Fig. 6). As the native *SCO3226* gene has been largely deleted in the complemented mutant the AbsA2 chIP peaks are attributable to protein binding to the SCO3226 DNA in the heterologous *3xF*-*absA2* construct expressed from pMT3226 integrated into the ϕC31*attP* site; furthermore the detected *absA2* transcripts are derived via transcription from the adjacent tandem *gylP1/P2* promoters in the expression vector. It is tempting to interpret these binding events in terms of a negative autoregulatory mechanism as AbsA2 has been shown to negatively regulate its own transcription, although an indirect mechanism where AbsA2 regulates its own expression *via* an intermediary, *e.g*. CdaR, could not be excluded [16]. We also note that the present study provides no evidence for AbsA2 binding to the promoter upstream of *absA1* (*SCO3225*) and that Sheeler *et al*., [17] also did not observe binding of AbsA2 to the *SCO3225* promoter region *in vitro*.

**Fig. 6.**
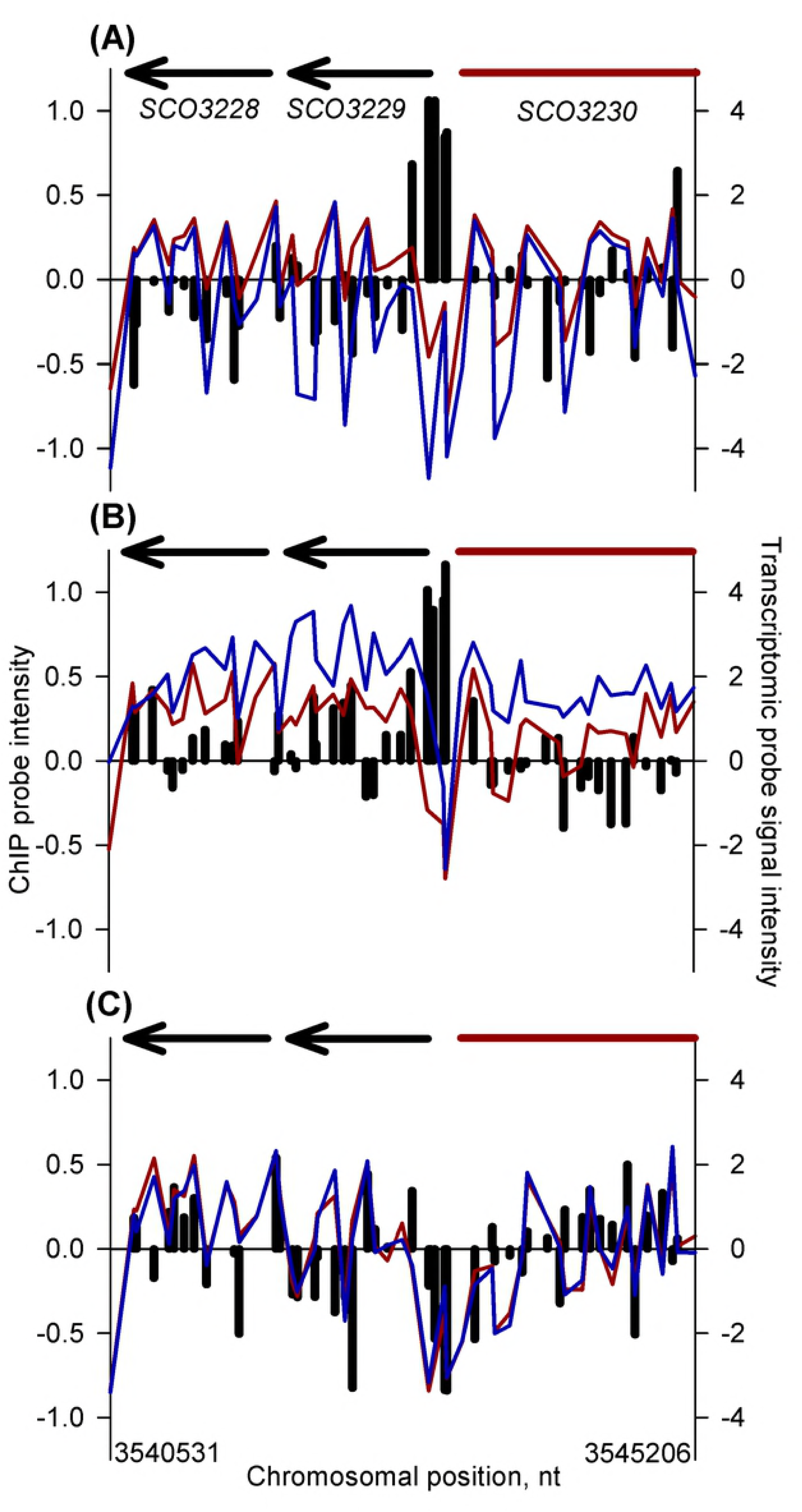
*In vivo* genomic distribution of 3xFAbsA2 in the region of *absA2* (*SCO3226)* and parallel measurement of adjacent gene expression.

See legend to Fig. 3 for an explanation of the axes and colour coding. Panels (A) & (B) refer to the 14 h and 18 h time-points, respectively.

The most interesting aspect of the *absA2* ChIP peak is that, in contrast to the ‘monophasic’ peak pattern observed upstream of *glmT*, *cdaPSI* and *cdaR,* the *absA2* ChIP peak is located *internal* to the gene and exhibits a curious asymmetrical, biphasic, form with a larger peak closest to the start of the gene overlapping with or followed by a smaller peak near the end of the gene.

### *In vivo* binding of AbsA2 to the pathway specific activator-encoding genes for RED and ACT, *redZ* and *actII*-*orfIV*

As expected from the literature [21], AbsA2 binds strongly to *redZ* at the mid-exponential growth phase, and although the AbsA2 ChIP peak persists into the 18 h late exponential phase time-point in the *3xFabsA2* strain, here it does not satisfy the ChIP enrichment scoring criteria (Fig. 7). The *redZ* AbsA2 ChIP biphasic peak pattern resembles that seen targeting *absA2* (*SCO3226*), with one large peak upstream of the gene and the second smaller peak within the gene. The fact that this unusual pattern is repeated at both the 14 h and 18 h time-points suggests that this is a genuine phenomenon and not an experimental artefact.

**Fig. 7.**
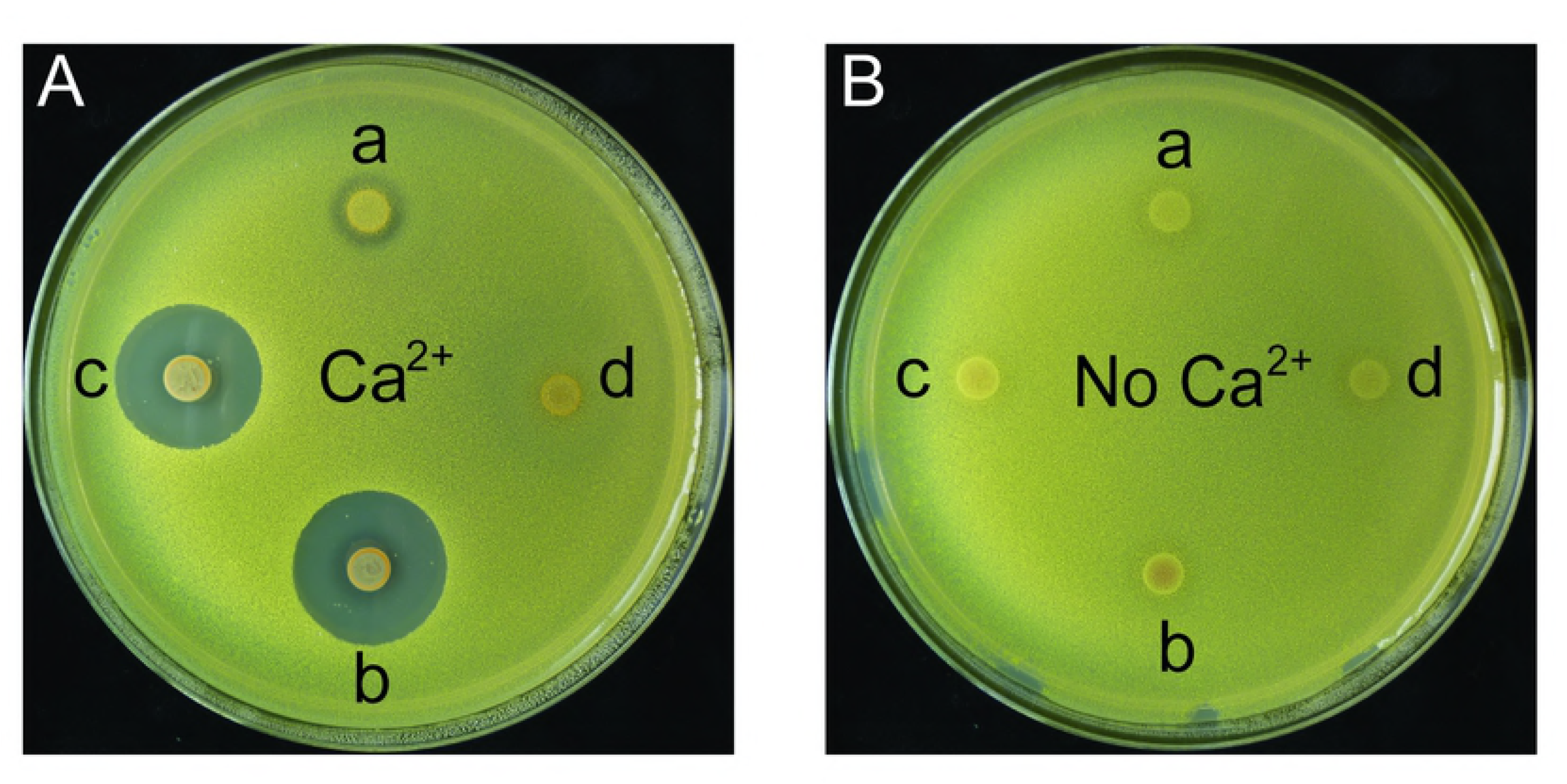
*In vivo* genomic distribution of 3xFAbsA2 in the region of *redZ* (*SCO5881)* and parallel measurement of adjacent gene expression.

See legend to Fig. 3 for an explanation of the axes and colour coding. Panels (A), (B) & (C) refer to the 14 h, 18 h and 35 h time-points, respectively.

A biphasic AbsA2 ChIP pattern is also observed in its binding to *actII*-*orfIV* (*SCO5085*) where AbsA2 only binds in the *3xFabsA2* strain sample at the mid-exponential time-point (Fig. 8). The lack of binding of AbsA2 at the later time-points correlates with a lifting of the repressive effect of AbsA2 which is indicated by the increase in expression of *actII*-*orfIV*, observed markedly in the stationary phase sample. It is noteworthy that although AbsA2 only binds to and represses *actII*-*orfIV* at the mid exponential time-point actinorhodin was not synthesized by the *3xFabsA2* strain until the culture had reached stationary phase. The regulation of actinorhodin biosynthesis has been described as *“amazingly complex”* [10], with a large number of transcription factors, including. AtrA [41], regulating ACT production. The effect on ACT biosynthesis observed here may therefore be due to long-lived, indirect, effects of AbsA2 exerting an influence on ACT biosynthesis until stationary phase, through disruption of a regulatory network of transcription factors long after the initial direct repression due to AbsA2 binding has been lifted.

**Fig. 8.**
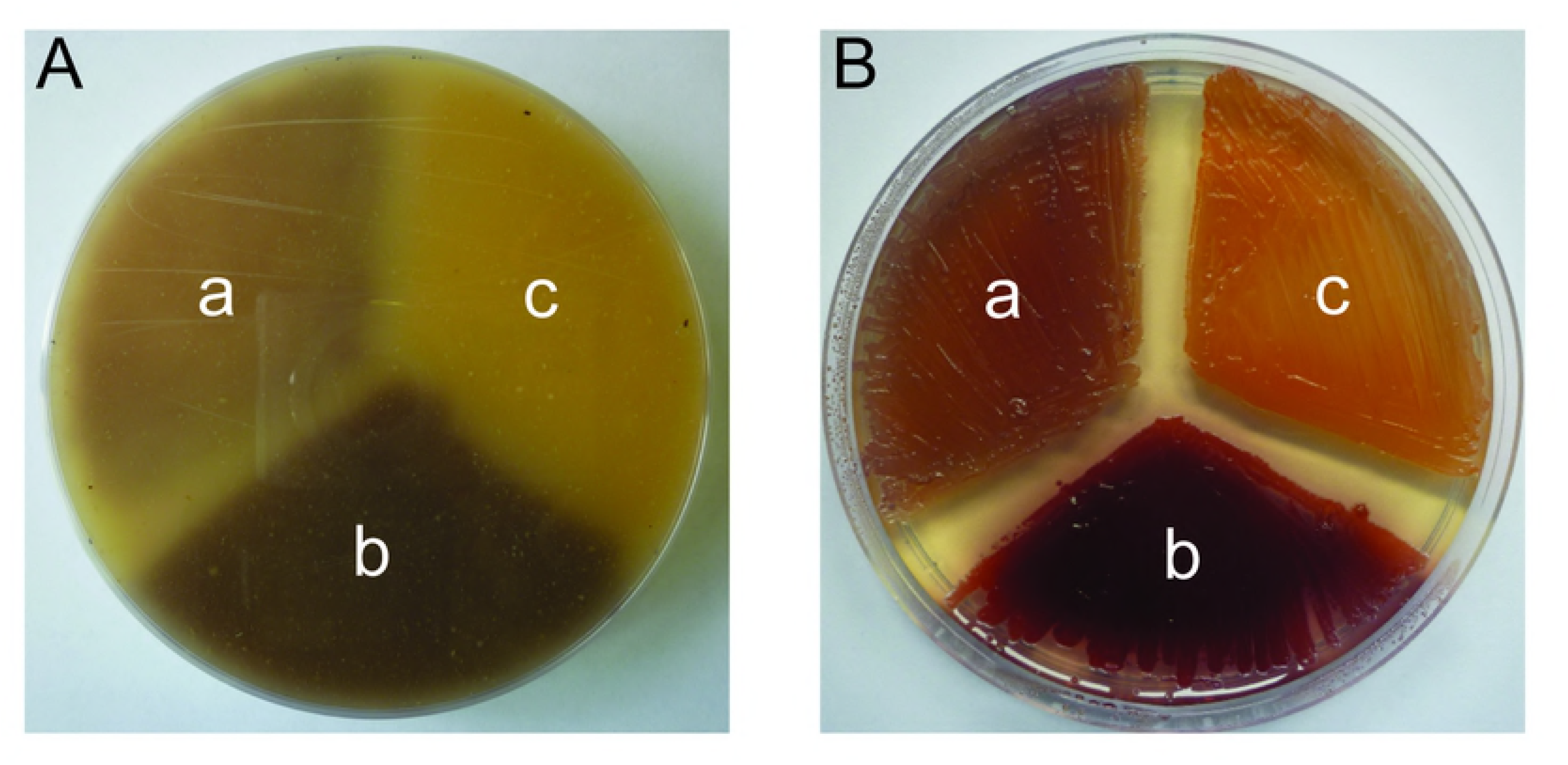
*In vivo* genomic distribution of 3xFAbsA2 in the region of *actII orfIV* (*SCO5085)* and parallel measurement of adjacent gene expression.

See legend to Fig. 3 for an explanation of the axes and colour coding. Panels (A), (B) & (C) refer to the 14 h, 18 h and 35 h time-points, respectively.

A number of regulatory mechanisms could potentially explain the biphasic AbsA2 binding patterns at *absA2, redZ* and *actII*-*orfIV*: AbsA2 binding to two sites independently; initially binding one site and then sliding along the DNA to the other site; AbsA2 binding at one point and then, following looping/coiling of the chromosome, contacting a second region of DNA at a remote location. As it has been shown that AbsA2 is capable of binding to short probes in gel-shift assays derived from the intergenic and 5′ terminus of *actII*-*orfIV* [21] it is unlikely that the double peak represents an allosteric binding event where the binding of AbsA2 to the upstream site is conditional on its occupancy of the downstream binding site.

It is tempting to interpret the differences in the mechanism of action of AbsA2 at (a) the *cdaR* and *cdaPSI* promoters and (b) the *redZ/actII*-*orfIV/absA2* genes in terms of the respective morphologies of the ChIP peaks, where AbsA2 associated with monophasic ChIP peaks exerts an indirect repressive effect on transcription by inhibiting activating transcription factors (e.g. CdaR), whereas the biphasic pattern of AbsA2 binding is associated with direct repression by AbsA2. Indeed, the fact that AbsA2 binding covers large portions of *redZ*, *actII*-*orfIV* and *absA2*, would suggest that the genes are more tightly constrained by AbsA2 occupancy and therefore less accessible to other transcription factors, than those with a single binding site.

We also note that the differences in AbsA2 ChIP peak form at the *redZ*, *actII*-*orfIV* and *cdaR* promoters correlate with the fact that AbsA2 has been shown to bind more weakly to the *cdaR* promoter than to the *redZ* and *actII*-*orfIV* promoters [21]. Our observed differences of *in vivo* AbsA2 binding patterns at the target loci also perhaps explain why previous investigators [21] have been unable to identify an AbsA2 consensus binding sequence and, despite our more detailed knowledge of the AbsA2 target sites, we have also been unable to identify consensus binding motifs. A possible explanation for this apparent lack of sequence specificity is that the binding of AbsA2 is dependent on factors other than the precise sequence of the target DNA *i*.*e*. in the case of *redZ* and *actII*-*orfIV* the double-peak binding pattern may be due to the effect of at least two widely spaced binding motifs, and possibly long-range DNA topological factors which influence DNA coiling/looping. In the case of the *cda* genes the presence/absence of binding DNA proteins (*e*.*g*. CdaR) may determine where AbsA2 binds. Indeed, it has been suggested previously that AbsA2 binding is dependent on the presence of CdaR [17].

The similarity of the *absA2* and the *redZ* and *actII*-*orfIV* AbsA2 ChIP peaks also suggests an interesting hypothesis as to how AbsA2 acquired the status of “master regulator” of antibiotic synthesis. It is possible that initially AbsA2 was merely a ‘pathway specific’ regulator of the *cda* biosynthetic gene cluster, in which it is located. However, due to the similarity in sequences of the *absA2*, *redZ* and *actII*-*orfIV* genes when, through horizontal gene transfer, the biosynthetic gene clusters were brought together in *S*. *coelicolor*, AbsA2 was able to bind the SARP genes of the ACT and RED clusters and so extend its control over their gene clusters, in addition to the *cda* cluster. This hypothesis is consistent with the fact that within the *cda* cluster AbsA2 operates as an integral part of a regulatory system in conjunction with the activator, CdaR, whilst due to it being a recently acquired regulator of *redZ* and *actII*-*orfIV* it acts independently at these loci, reflecting its recent integration into the endogenous transcription factor network.

## Conclusion

The present study used the powerful tool of high-density microarrays to investigate the regulation of antibiotic production in *Streptomyces coelicolor* by the transcriptional regulator AbsA2. Despite myriad genetic and molecular biological studies of this transcription factor the present study represents the first attempt to integrate a transcriptomic study with ChIP-on-chip technology. The results indicate that AbsA2 binds to several loci within the *cda* cluster (*SCO3215*, *SCO3217*, and *SCO3230*) and suggest that it exerts a repressive influence on transcription through negatively regulating CdaR activity. Additionally, we show that AbsA2 also binds to its own gene (*SCO3226*), providing evidence for an autoregulatory mechanism, and to the SARP genes of the *act* (*actII*-*orfIV*) and *red* (*redZ*) clusters; differences between the patterns of the AbsA2 ChIP peaks at these genes and those within the *cda* cluster suggest that AbsA2 represses transcription of these regulatory genes directly and raises the intriguing possibility that AbsA2 displays two different mechanisms of binding DNA and of mediating repression.

The results of the study go some way towards explaining and illuminating aspects of previous genetic and molecular studies of AbsA2. They also raise new questions as to the precise mode of action of AbsA2 and it is hoped that these results will serve to inform and stimulate future work into this fascinating transcription factor.

## Supplementary Figure Legends

**Fig. S1. DNA sequence of the N-terminally triple flag tagged *absA2* and upstream flanking region**.

The green sequences represent restriction enzyme cut sites incorporated into the termini for ease of cloning. The black sequence represents the *absA1*-*absA2* intergenic region, the red sequence represents the DNA which encodes the triple-flag tag and the blue sequence represents the *absA2* coding sequence.

**Fig. S2. Pigmented antibiotic production in liquid medium**.

Actinorhodin (ACT) and undecylprodiginine (RED) assays from replicate cultures of strains of MT1110 *ΔabsA2* (pMT3226) and MT1110 *ΔabsA2* (pMT3226::*3xFabsA2*).

**Fig. S3. Results of the transcriptomic study**.

List of genes which are significantly differentially expressed between MT1110 *ΔabsA2* (pMT3226) and MT1110 *ΔabsA2* (pMT3226::*3xFabsA*) at the 14 h, 18 h and 35 h time-points. Genes in red are repressed in MT1110 *ΔabsA2* (pMT3226::*3xFabsA*) relative to MT1110 *ΔabsA2* (pMT3226). Genes highlighted in green are activated in MT1110 *ΔabsA2* (pMT3226::*3xFabsA*) relative to MT1110 *ΔabsA2* (pMT3226). Genes which possess clusters of significantly ChIP-enriched microarray probes in MT1110 *ΔabsA2* (pMT3226::*3xFabsA*) relative to MT1110 *ΔabsA2* (pMT3226) are highlighted by green/red background shading. Genes of the *act* biosynthetic gene cluster are highlighted by red borders and genes of the *cda* biosynthetic gene cluster are highlighted by blue borders.

## Authors Contributions

RAL constructed the complemented *absA2* mutant strain, did the ChIP-on-chip and transcriptomic experiments, analysed the microarray data and wrote the manuscript. AW constructed and characterised the *ΔabsA2* mutant strain. GB assisted with the microarray experiments and manuscript preparation. EEL processed the transcriptomic microarray data. CM-L processed the ChIP-on-chip microarray data. AK led the project. CPS led and managed the project, designed the experiments and contributed to the manuscript.

